# A Novel GelMA-OrnMA Electrically Conductive Bioink for Developing Engineered Neural Tissues

**DOI:** 10.1101/2024.10.06.616594

**Authors:** Mahmoud A. Sakr, Kartikeya Dixit, Kinam Hyun, Sumi Siddiqua, Su Ryon Shin, Hitendra Kumar, Keekyoung Kim

## Abstract

Electrical conductivity is a crucial requirement of matrices for developing engineered neural tissues. A conductive matrix not only supports cell growth but also provides potential to stimulate the cells. However, electrically conductive matrices often require inclusion of synthetic polymers, nanomaterials and large number of ionic species. While enhancing electrical conductivity, often properties like transparency, mechanical stiffness and biocompatibility are compromised which can render the resulting matrices partially suitable for neural tissue engineering. Further, the byproducts of matrix degradation can have unforeseen influences. Therefore, electrically active matrices are required which provide a suitable combination of electrical conductivity, mechanical properties and biocompatibility. In this work, a novel biomaterial is described which results in optically transparent, electrically conductive and highly biocompatible matrices along with ability to match the native neural tissue stiffness. Using gelatin methacryloyl (GelMA) as base hydrogel, we covalently incorporated zwitterionic functional groups to obtain a composite matrix. The zwitterion moieties were derived from Ornithine by synthesizing ornithine methacryloyl (OrnMA) and blending with GelMA inks. Through systematic characterization we demonstrated the suitability of GelMA-OrnMA hydrogels in providing mechanical stiffness matching the native neural tissues, supporting proliferation of human astrocytes in 3D culture and electrical conductivity in the range required for electrically active cell types like astrocytes. Owing to their electrical conductivity, these matrices also influenced the growth of astrocytes which manifested as significant changes in their organization and morphology. These findings suggest that GelMA-OrnMA has immense potential as a bioink for developing engineered neural tissues.

## 1. Introduction

Hydrogels are the most used biomaterials in tissue engineering^1–3^. To advance healthcare, biomaterials research is expanding towards developing advanced materials that integrate functionalities ^2,4–7^. Among these advanced materials are electrically conductive hydrogels, which are frequently used in tissue engineering applications due to their unique physical and electrical properties that promote functional tissues^8–10^. There are various methods for synthesizing conductive hydrogels to achieve biocompatibility and restore cellular functions^11–13^. Metal nanoparticles are commonly used in these hydrogels^14–16^. It is reported in the literature the use of metal nanoparticles and combining them with hydrogels is for the sake of adding new functionalities to the base hydrogel. Gold, silver, and platinum nanoparticles are great candidates because of their antimicrobial properties and their electrical conductivity^17^. As mentioned in Kiyotake et al.^18^, they reported the use of gold nanorods with hyaluronic acid and gelatin hydrogel. Their study demonstrated the fabrication of non-toxic citrate-gold nanoparticles. This study explained the effect of gold nanoparticles in supporting the adhesion and viability of rat neural stem cells. Electrically conductive functionalized hydrogels with carbon nanotubes (CNTs) and graphene oxide have a potential for stimulating nerve cell differentiation^19^. It is reported that CNTs reinforce the stiffness of the hydrogel without affecting the porosity of the hydrogel^20^. The disadvantage of using CNTs and graphene oxide is their aggregations within the hydrogel because of their hydrophobicity. These aggregates affect the transparency of the material^21^. Polypyrole (PPY) is a widely used and studied conductive polymer in neural development^22^. It is mentioned that PPY can enhance cell adhesion, proliferation, and differentiation with or without electrical stimulation^23^. Poly(3,4-ethylenedioxythiophene) (PEDOT) and polyaniline (PANI)are other conductive polymers that exist in literature and are used in tissue engineering^24^.

Zwitterionic polymers are a type of polymer that contain an equal number of cations and anions in their polymer chain, providing several beneficial properties such as antifouling, functionalization, and flexibility of design^25,26^. The cations and anions in the macromolecular chains of zwitterionic polymers make them hydrophilic, and they exhibit anti-polyelectrolyte behaviour by enlarging in aqueous solutions with the presence of ions^27^. These characteristics make zwitterionic polymers suitable for tissue engineering applications, and zwitterionic hydrogels have several advantages over other hydrogel materials used in 3D bioprinting, such as low non-specific protein adsorption^28^, high swelling ratio, and tunable mechanical properties that can mimic different tissues in the body^29^.

Ornithine is a naturally occurring amino acid with several physiological functions, such as enhancing wound healing, antioxidant, anti-inflammatory, and detoxification of ammonia in the liver^30–32^. Ornithine has been integrated with hydrogels to work as a biomaterial for tissue engineering applications^33–36^. In addition, ornithine induces neuronal differentiation from neural/progenitor stem cells and induces biological behaviors such as cell growth, proliferation, and differentiation^37^. The addition of ornithine to hydrogels creates a porous and biodegradable structure that has shown promising results in tissue engineering and regenerative medicine^38^.

This article discusses the development of a new bioink that is electrically conductive and can be used for neural tissue engineering applications. Ornithine shows a promising effect with gelatin methacryloyl (GelMA) regarding its electrical conductivity and biocompatibility. It is also noticed that ornithine enhances the mechanical stiffness of pristine GelMA and can modify the pore size and the swelling property of the bio-ink. Astrocytes maintained their proliferation and elongation along the surface and within the ink matrix.

## 2. Materials and methods

### 2.1. Ornithine methacrylate synthesis

Ornithine methacryloyl (OrnMA) was synthesized by reacting L-Ornithine (Sigma Aldrich, St. Louis, MO, USA) with glycidyl methacrylate (GMA) (Sigma Aldrich, St. Louis, MO, USA). First, 2 g of L-Ornithine was dissolved in 20 mL of distilled water at room temperature. Next, 4 mL of GMA was added dropwise to the L-Ornithine solution while continuously stirring at 500 rpm. The reaction was continued for 12 hours or 24 hours in dark with continuous stirring at 500 rpm. Next, 60 mL ethanol was cooled at −20°C for 15 minutes. Then the L-Ornithine solution was added dropwise to the cold ethanol while continuously stirring at 200 rpm. After complete addition, the resulting mixture was left undisturbed for 10 minutes. Next, the mixture was agitated and transferred to 50 mL tubes and centrifuged to collect the precipitate. The supernatant was decanted, and 20 mL of fresh ethanol was added to each tube, followed by vigorous agitation and centrifugation. This process was repeated three times to obtain a filtered precipitate of OrnMA. Next, the precipitate was dissolved in 20 mL distilled water. The resulting OrnMA solution was frozen at −80 °C for 12 hours and then lyophilized for 3 days. The OrnMA powder obtained after lyophilization was stored at −20 °C for further usage. Methacryloyl substitution on ornithine was confirmed using as reported previously ^39–41^. Concentrated solutions of ornithine and OrnMA were prepared by dissolving 0.8 g of respective materials in 0.6 mL of deuterium oxide (Acros Organics, Fisher scientific). A 400 MHz spectrometer (Bruker, Billerica, MA, USA) was used to record the ^1^HNMR spectra for each of the samples. The recorded data was preprocessed by performing phase correction, suitable baseline fitting and scaling in the MestReNova software (Mestrelab Research, Santiago de Compostela, Spain).

### 2.2. GelMA-OrnMA prepolymer inks preparation

Gelatin methacryloyl (GelMA) was synthesized by reacting gelatin with GMA following a protocol outlined previously with some modifications^42^. In brief, 5 g of gelatin from porcine skin (Type A, bloom strength 300, Sigma Aldrich, St. Louis, MO, USA) was dissolved in 50 mL distilled water heated at 50°C. Continuous stirring at 500 rpm was maintained throughout the synthesis process. Once a homogenous solution was formed, 10 mL of GMA was added dropwise. The resulting solution was then left undisturbed at 50°C and 500 rpm for 12 hours in the dark to continue the reaction. After 12 hrs, 50 mL of distilled water was added to the solution to stop the reaction. The final solution was then transferred to dialysis tubing (12-14 kDa, Fisher Scientific) for dialysis against distilled water. The dialysis was performed at room temperature, and the dialysis water was changed twice a day for 3 days. After dialysis process, the resulting GelMA solution was transferred to 50 mL tubes and frozen at −80°C for 24 hours. Next, the GelMA samples were lyophilized in a freeze dryer for 3 days to obtain a dried foam. The lyophilized GelMA was stored at −80°C for further use.

The GelMA-OrnMA prepolymer inks were prepared by first dissolving GelMA in PBS at 5% (w/v) concentration. Next, OrnMA powder was added to the GelMA solution up to the concentrations of 5%, 10% and 15% (w/v). Finally, the two component photoinitiator comprising 0.02 mM Eosin Y (EY) and 0.2% (w/v) triethanolamine (TEOA). The complete solution was agitated for mixing all the components. Table 1 lists the compositions of various prepolymer inks.

**Table 1.** GelMA-OrnMA ink compositions.

| Macromer |  | Photoinitiator |  | Prepolymer ink<br>code |
| --- | --- | --- | --- | --- |
| GelMA | OrnMA | Eosin Y | Triethanolamine |  |
| 5% | 0 | 0.02 mM | 0.2% | 5% GelMA |
|  | 5% |  |  | 5% GelMA-5% OrnMA |
|  | 10% |  |  | 5% GelMA-10% OrnMA |
|  | 15% |  |  | 5% GelMA-15% OrnMA |

### 2.3. Photocrosslinking characterization

First, the optical transparency of the GelMA-OrnMA inks was characterized. Different compositions of GelMA-OrnMA inks were prepared by sequentially dissolving GelMA and OrnMA in PBS. Photoinitiator components were not added to these solutions. After complete dissolution, 1 mL prepolymer ink was dispensed in a 1 cm cuvette and placed in a real-time UV-Vis spectroscope (FLAME-T-UV-VIS, Ocean Optics, FL, USA). An in-built illumination source of the spectroscope was used and the transmission spectra for various ink compositions were recorded. Transmission spectrum of PBS was also recorded as reference.

The photocrosslinking characteristics of the GelMA-OrnMA inks were evaluated by recording the variation in the rheological properties with visible light exposure. The rheological characterizations were conducted using an MCR 302e rheometer (Anton Paar, Austria). A stainless-steel parallel plate with a diameter of 25 mm was used as a measuring set. During the rheological measurement, a gap distance of 0.5 mm was maintained between the probe plate and the rheometer platform. All measurements were performed at room temperature. To record the photocrosslinking kinetics, a transparent glass plate was integrated into the setup, allowing for the illumination of the hydrogel with visible light from the bottom of the measuring set. After initial probing of the shear modulus (storage (G’) and loss modulus (G”)) for 1 minute, the visible light irradiation was initiated and continued for a duration of 5 minutes. After 5 minutes, the light source was turned off and subsequent rheological measurements were continued. Further, oscillatory measurements were conducted with a 1% shear strain and a frequency of 1 Hz, and both the G’ and G” for the photocrosslinked hydrogels were simultaneously recorded.

### 2.4. Microstructure analysis

The porous structure of hydrogels plays a crucial role in determining their physical characteristics as well as survival of the cells encapsulated within the hydrogel matrix. Therefore, the microscale porous structure of the hydrogels was visualized. First, the hydrogel prepolymer solutions were prepared as described earlier with compositions listed in Table 1. Next, the prepolymer solutions were dispensed to fill a cylindrical mold of 8mm diameter and 8mm height. Next, the hydrogels were formed by photocrosslinked using visible light exposure for a duration of 10 minutes to ensure complete photocrosslinking. The formed hydrogels were retrieved and immersed in distilled water for 24 hours to achieve equilibrium swelling. Next, the hydrogels were frozen at - 80°C and then freeze dried for 3 days. The dried samples were carefully broken using a sharp blade to expose the cross-section and then loaded on metal stubs with carbon tape. The samples were then sputter coated with a 10 nm layer of Au/Pd alloy. Next, a field emission scanning electron microscope FESEM (MIRA3 XMU, TESCAN, Brno, Czech Republic) was used to acquire images to visualize the porous structure of the dried hydrogels. The pore size and pore distribution along the micro walls of the freeze-dried different concentrations of ornithine with 5% GelMA were visualized and compared in the obtained images. A custom MATLAB script was used for average pore size calculation and pore size distribution.

### 2.5. Mechanical and swelling behavior characterization

For characterizing the mechanical properties of the GelMA-OrnMA hydrogels, cylindrical samples were fabricated by dispensing pre-polymer solution in molds with 8 mm diameter and 8 mm height. The crosslinked hydrogels were formed upon exposure to visible light (48.7 mW/cm^2^) in a custom-made bioprinting system^42^. The exposure duration was kept 10 minutes for ensuring complete crosslinking of the large volume and thickness of the pre-polymer solution layer. After photocrosslinking, the samples were carefully retrieved from the molds and immersed in PBS overnight to achieve equilibrium swelling. The Young’s modulus of GelMA-OrnMA hydrogel samples was measured using a mechanical testing platform (MACH-1, Biomomentum, Laval, QC, Canada). Prior to measurement, the samples were blotted to remove excess PBS on the surface. A maximum strain of 50% was applied on the samples using a 12mm diameter flat probe. The force-displacement data for each specimen was recorded and analyze using a custom-made MATLAB script. The stress-strain data was obtained from force-displacement data by using the physical dimensions of the hydrogel samples before compression. Next, the slope of the linear region (approximately up to 10% strain) was used to compute Young’s modulus of the hydrogel samples. The crosslinked GelMA-OrnMA hydrogels were prepared for examining the swelling behavior following a similar hydrogel preparation method as described above. After photocrosslinking, the hydrogel samples were immersed in PBS overnight for achieving equilibrium swelling. Next, the samples were gently blotted to remove excess liquid on the surface and the weight of the swollen hydrogel samples was recorded. Next, the hydrogels were transferred to a 24-well plate and frozen at −20°C for 12 hours followed by lyophilization for 3 days. Further, the weight of the dried specimens was recorded and compared with the weight of corresponding hydrated specimens recorded earlier. The swelling ratio depicting water uptake capacity of the hydrogels was calculated using equation Eq. 1.

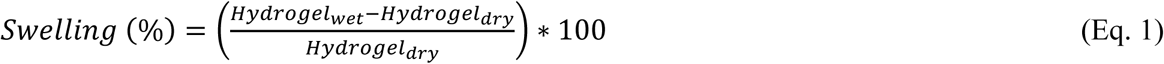

### 2.6. Electrical conductivity measurements

The electrical conductivity of the hydrogels was measured using electrochemical impedance spectroscopy (EIS, PalmSens4, PalmSens BV, Netherlands). In brief, hydrogel samples were prepared with 5% w/v GelMA and 0%, 5%, 10%, and 15% w/v OrnMA, respectively. The photocrosslinking duration was fixed at 3 minutes for all samples, and the dimensions of the hydrogels were maintained at 24 mm × 24 mm × 2 mm (L×W×H). Additionally, two sets of these samples were prepared using distilled water and PBS as solvents. A custom 3D-printed platform was fabricated to place the samples between electrodes made of glass slides covered with conductive copper tape. After crosslinking, the hydrogel samples were wiped to remove excess liquid and placed between two electrodes. The ends of the electrodes were connected to the EIS station.

The conductivity measurements were performed at room temperature with a frequency range from 0.1 Hz to 1 MHz, with an applied potential of 10 mV and 1 mV amplitude. Data were acquired and analyzed with PSTrace software (PalmSens BV, Netherlands). The conductivity of the hydrogels (σ) was calculated using Eq. 2, where (*l*) represents the distance between the electrodes and (*A*) represents the area of the electrode in contact with the sandwiched hydrogel sample. The resistance of the hydrogel samples (*R*) was determined by fitting the Nyquist curve with a Randles equivalent circuit model^43,44^.

The conductivity measurements were performed at room temperature with a frequency range from 0.1 Hz to 1 MHz, with an applied potential of 10 mV and 1 mV amplitude. Data were acquired and analyzed with PSTrace software (PalmSens BV, Netherlands). The conductivity of the hydrogels (σ) was calculated using Eq. 2, where (*l*) represents the distance between the electrodes and (*A*) represents the area of the electrode in contact with the sandwiched hydrogel sample. The resistance of the hydrogel samples (*R*) was determined by fitting the Nyquist curve with a Randles equivalent circuit model^43,44^.

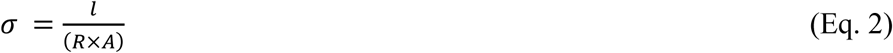

### 2.7. Biocompatibility assessment of the hydrogels

The U118-MG human astrocytes (HTB-15, ATCC, Manassas, VA, USA)) were maintained in culture at 37°C and 5% CO_2_ atmosphere and using a growth media comprising Dulbecco’s Modified Eagle Medium (DMEM) (Lonza, Basel, Switzerland) supplemented with 10% fetal bovine serum (FBS) and 1% penicillin-streptomycin. For subsequent experiments, the cells were first treated with trypsin-EDTA solution to detach from the cell culture flasks. Next the media with cell suspension was centrifuged to collect the cells in the form of a pellet. Further, the pellet was resuspended in respective bioinks to obtain desired cell density.

The astrocytes were, first, in a 3D culture *in vitro* model by encapsulating them in GelMA-OrnMA hydrogels. A 500 µm thick layer of GelMA-OrnMA was photocrosslinked under the custom bioprinting system^42^ in a mold with a cell encapsulation density of 2 × 10^6^ cells/mL. After crosslinking, the samples were washed three times with PBS and then carefully transferred from the mold to a 12-well plate. Each well was filled with 3 mL of cell growth media to completely immerse the samples. The samples were maintained in culture at 37°C and 5% CO_2_ atmosphere and growth media was replenished every two days. The brightfield images of the samples were recorded over a span of 7 days to observe the behavior of the encapsulated cells using an inverted fluorescence microscope (Axio Observer 7, Carl Zeiss Canada Ltd, North York, ON, Canada).

The biocompatibility characterization of the cell-laden hydrogels was done by using a standard live/dead assay (Biotium, Fermont, CA, USA). The assay comprised fluorescent dyes Calcein-AM for staining live cells and ethidium homodimer III (EthD-III) for staining dead cells. The preparation of a working staining solution of 2 µM Calcein-AM and 4 µM EthD-III included addition of 2 µL Calcein-AM (4 mM in DMSO stock) and 10 µL EthD-III (2 mM in DMSO/H_2_O stock) in 5 ml of sterilized PBS. The hydrogel samples with encapsulated cells were washed twice with PBS and stained with the working solution for 20 mins at 37 °C in dark conditions. The samples were washed twice with PBS after staining and then imaged under an inverted fluorescence microscope (Revolve, ECHO, San Diego, CA, US). The FITC and Texas Red channels were used to visualize and distinguish live and dead cells, respectively and then merged to form the representative images.

The proliferation of the astrocytes encapsulated in GelMA-OrnMA hydrogels was assessed using a metabolic activity assay. A colorimetric assay using XTT (sodium 3’-[1-(phenylaminocarbonyl)-3,4-tetrazolium]-bis (4-methoxy6-nitro) benzene sulfonic acid hydrate) was used in the form of a two-reagent kit (XTT, Biotium, Fermont, CA, USA). The working solution was prepared immediately before use as per manufacturer’s instructions by mixing 25 µL Activation Reagent with 5 mL XTT Solution. The cell encapsulating hydrogel disk samples (diameter 6 mm and thickness 1mm) were transferred to a 24-well plate and incubated with 100 µL activated XTT solution and 200 µL cell growth media for 6 hours at 37 °C. After incubation, 100 µL supernatant from each well was collected loaded on a 96-well plate. The 96-well plate was then examined in a microplate reader (Bio-Rad, Philadelphia, USA) to record absorbance at 490 nm wavelength.

### 2.8. Cell morphology characterization

The U118 human astrocytes maintained in culture were collected by trypsinization followed by centrifuging. GelMA-OrnMA bioinks were prepared using the compositions listed in Table 1 and the collected cells were added to the bioinks to achieve a density of 2 × 10^6^ cells/mL. The bioinks were photocrosslinked to form cell encapsulating hydrogels with a thickness of 500 µm. The hydrogel samples were maintained in culture using the cell growth media for further examination. After culturing for 7 days, the samples were fixed using 4% paraformaldehyde for 15 minutes. Next, the samples were washed twice with PBS and then incubated with 0.1% Triton X-100 solution in PBS at room temperature for 10 minutes. The samples were washed again with PBS for three times. Next, the fixed hydrogel samples were incubated with 100 nM working solution of Phalloidin (Acti-stain 488, Cytoskeleton, Denver, CO, USA) for 60 minutes at room temperature. The samples were washed thrice with PBS and then mounted using a mounting media containing 100 nM 4’,6-diamidino-2-phenylindole (DAPI) (Fluoroshield with DAPI, Sigma-Aldrich) for counterstaining DNA. Next, the samples were imaged using an inverted fluorescence microscope (Axio Observer 7, Carl Zeiss Canada Ltd, North York, ON, Canada). DAPI and FITC channels were used to capture images corresponding to stained nuclei with DAPI and stained cytoskeleton with phalloidin respectively. The morphology of the cells in various hydrogels was examine using the captured DAPI/Phalloidin staining images. The image area covered by the cells was measured as one of the markers by calculating the number of pixels. Another parameter measured was nuclei aspect ratio from the DAPI channel images.

### 2.9. Bioprinting

The U118 astrocytes maintained in culture were collected after trypsinization and centrifuging. Next, the cells were suspended in a 5% GelMA-10% OrnMA solution to obtain a bioink with 2 × 10^6^ cells/mL. A high-resolution scaffold was printed using a custom bioprinting system^42^. The scaffold featured thin branches connecting rings. After bioprinting, the structures were washed twice with PBS and then cultured in cell growth media for 5 days. The samples were visualized using brightfield imaging at days 0, 3 and 5 to examine the cell growth within the samples. On day 5, the samples were fixed and stained with DAPI and phalloidin as described above to visualize the cell organization within the scaffold matrix.

### 2.10. Statistical analysis

All data are presented as mean values accompanied by the standard deviation. Statistical analysis was conducted utilizing GraphPad. To determine statistical significance, a one-way analysis of variance was employed along with Tukey’s multiple comparison test. Differences were regarded as statistically significant when the p-value was less than 0.05, and the significance levels were indicated as follows: ns (not significant) for p > 0.05, * p < 0.05, ** p < 0.01, *** p < 0.001, and **** p < 0.0001.

## 3. Results and discussion

### 3.1. L-Ornithine methacryloyl synthesis

L-Ornithine was chemically modified to obtain L-Ornithine methacryloyl (OrnMA). This was achieved by substitution of methacryloyl functional groups on primary amine of ornithine. A highly simplified two-step synthesis process was designed in which the first step corresponds to the substitution reaction and the second step corresponds to the filtration and retrieval of the final product. Figure 1A elaborately depicts the sequence of steps involved in the two-step synthesis process. High solubility of L-Onithine in water led to the selection of distilled water as preferred solvent for the reaction. Further, glycidyl methacrylate (GMA) was selected as precursor for the reaction. This selection was guided by gelatin methacryloyl (GelMA) synthesis optimization reported earlier which demonstrated successful methacrylation of gelatin using GMA as precursor and distilled water as solvent^42^. The reaction was carried out with 2 g of L-Ornithine monohydrochloride (molecular weight: 168.62 g/mol) which translated to 11.86 mmol. In order to obtain a 1:2.25 molar ratio of ornithine to GMA, the amount of GMA (molecular weight: 142.15 g/mol, density: 1.07 g/mL) was kept 26.68 mmol which translated to approximately 4 mL. This ratio of ornithine to GMA was lower than typical 10x or 20x ratio used in GelMA synthesis^42,45,46^. This was done to limit the extent of substitution of methacryloyl functional groups on ornithine such that OrnMA zwitterions could be obtained. Since GMA is not miscible with distilled water, the solution was stirred continuously at 500 rpm throughout the reaction duration to form an emulsion comprising small droplets of GMA in aqueous phase. This allowed a larger contact surface area between the GMA and L-Ornithine dissolved in the aqueous phase for the substitution reaction to continue. After the synthesis continued for 12 hours, the resulting solution was added to ethanol. In this step, ethanol acted as a bridge solvent exhibiting miscibility with water as well as GMA. Therefore, all the liquid components formed a single phase. On the other hand, ornithine and ornithine methacryloyl (OrnMA) have lower solubility in ethanol as compared to water^47,48^. Further, this solubility difference can be increased by lowering the temperature. Therefore, ornithine and OrnMA were precipitated in a powdered form after adding the solution to cold ethanol. However, the precipitation was not instantaneous and requires up to 30 minutes. After the formation of the precipitate, the mixture was centrifuged to collect precipitate and supernatant was discarded. In order to ensure complete removal of GMA, the precipitate was resuspended in ethanol and agitated using a vortex mixer followed by centrifuging. This process was repeated twice and referred to as two washes by ethanol in this study. The collected OrnMA precipitate was then dissolved in distilled water and frozen at −80 °C. Finally, the frozen OrnMA solution was freeze dried to obtain OrnMA powder. The scheme of methacrylate substitution reaction during OrnMA synthesis is shown in Figure 1B. In order to confirm the methacrylate substitution, proton nuclear magnetic resonance (^1^HNMR) was used as reported previously ^39–41^. Figure 1C shows the respective ^1^HNMR spectra of L-Ornithine and OrnMA stacked for comparison. The presence of methacrylate vinyl signal (6.0–6.2 and 5.6–5.8 ppm) and the presence of a signal from the methyl group from the methacrylate functional group (1.2–1.8 ppm) indicated the successful substitution of methacrylate groups on ornithine to result in OrnMA.

**Figure 1.**
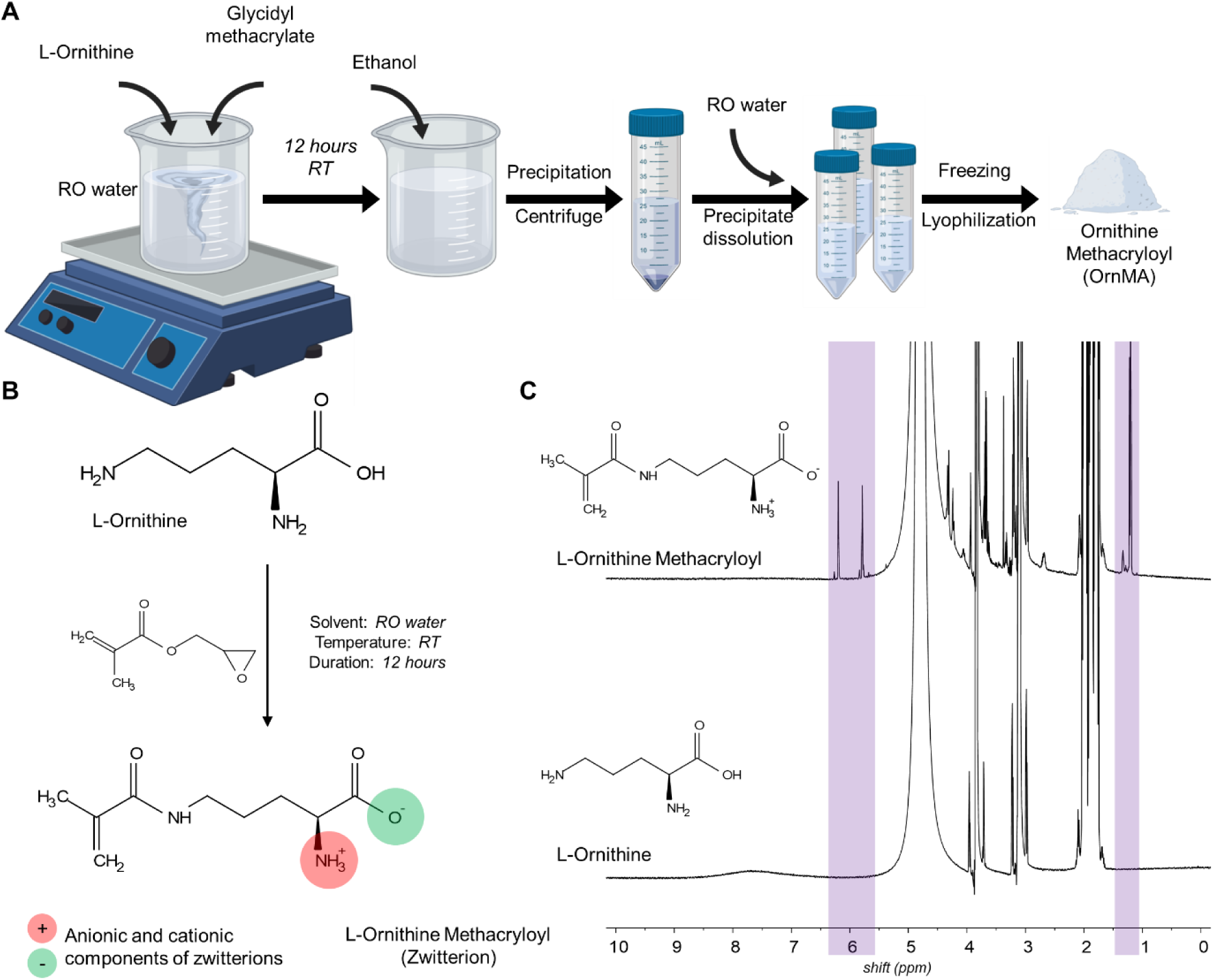
Ornithine methacryloyl (OrnMA) synthesis. (A) Schematic representation of the sequence of steps involved in OrnMA synthesis process. (B) Reaction scheme of methacrylate substitution on ornithine. (C) Comparison of ^1^HNMR spectra of L-ornithine and OrnMA.

Despite methacrylate functionalization, OrnMA polymerization cannot yield a stable hydrogel network capable of forming a defined shape and rather results in a polymerized ionic liquid. Therefore, GelMA was selected as a base biomaterial in this work and OrnMA was co-polymerized with GelMA to obtain composite hydrogels. GelMA was synthesized using a well-established protocol reported in earlier studies^42,49,50^. The GelMA obtained using this protocol has been reported to exhibit more than 80% degree of substitution and low gelation temperature to avoid gelation as room temperature^42^.

### 3.2. GelMA-OrnMA inks characterization

GelMA-OrnMA inks were prepared at various compositions as listed in Table 1 including pristine GelMA ink. First, the optical transparency of these inks was evaluated since it can significantly influence the intensity distribution of the transmitted light within the ink layer. A visual inspection (Figure 2A) showed visibly clear solutions of pristine GelMA, 5% GelMA-5% OrnMA and 5% GelMA-10% OrnMA compositions indicating high optical transparency. However, the 5% GelMA-15% OrnMA solution appeared slightly turbid. Further, visible light transmission through these inks was characterized using an UV-Vis spectrometer. For reference, the spectrum of the transmitted light through PBS was also recorded. During this process, the light intensity and spectrum capture parameters were fixed to allow comparison of relative transmitted light intensity. The composition with 5% GelMA-10% OrnMA exhibited highest optical transparency of approximately 95% among the set of inks followed by 5% GelMA-5% OrnMA ink with optical transparency of approximately 90%. Composition with 5% GelMA, on the other hand, exhibited an optical transparency of approximately 85%. The 5% GelMA-15% OrnMA solution, observed to be turbid visually, allowed approximately 80% transmission of incident light. Along with differences in the optical transmission from various inks, the minimum transparency observed was 80% which can potentially influence the photocrosslinking of these inks by introducing variations in the light intensity within the ink layer. Figure 2 A and B demonstrates the transparency of the GelMA-OrnMA, as shown in Figure 2 B, the transparency of the hydrogel is maintained while increasing OrnMA’s concentrations. This data demonstrates the ability of GelMA-OrnMA hydrogels for applications that require a transparent hydrogel such as skin tissues.

**Figure 2.**
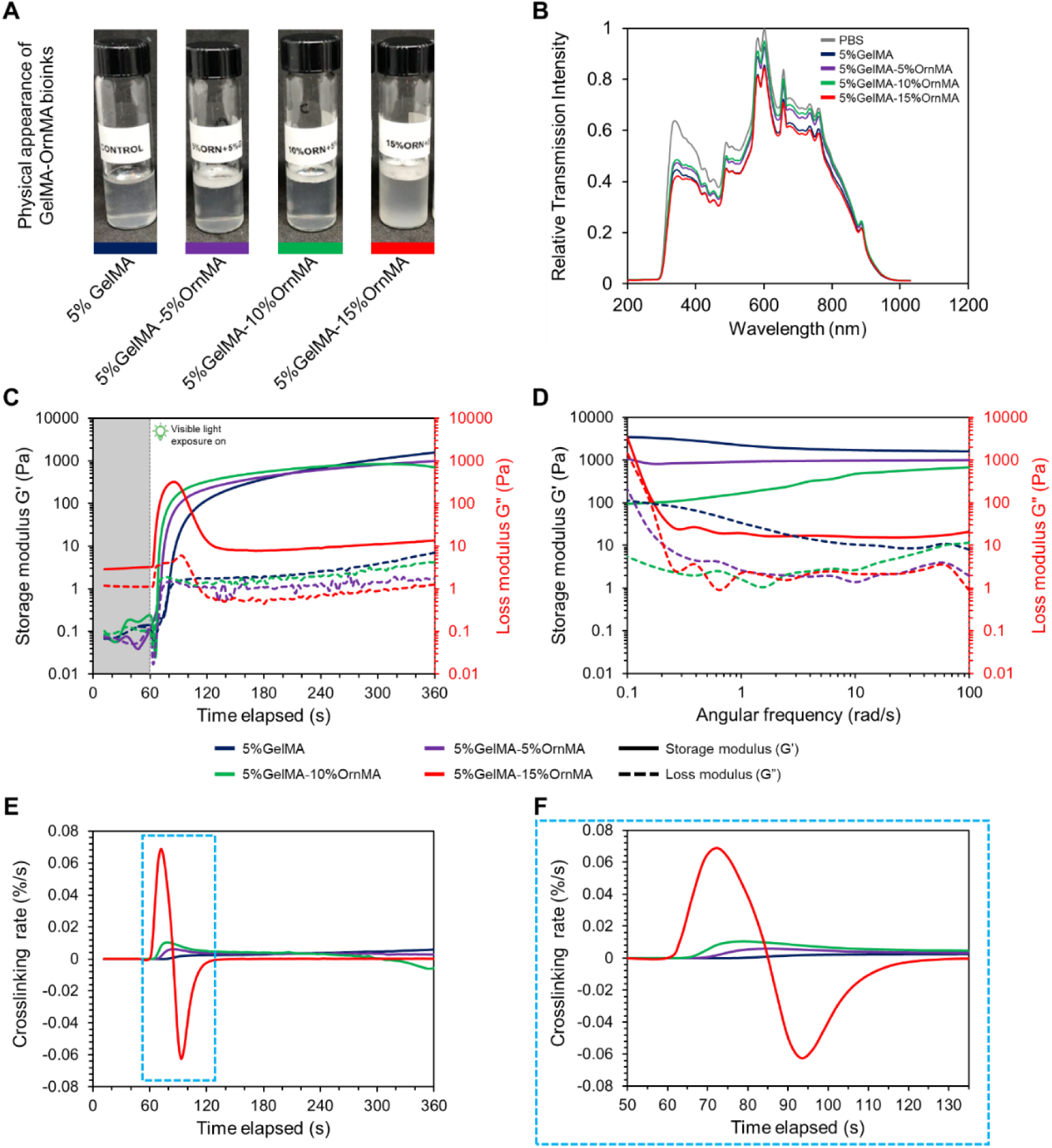
(A) Physical appearance of GelMA-OrnMA hydrogels. (B) Transparency evaluation using UV-Vis Spectrometry. (C) Photocrosslinking characterization of GelMA-OrnMA composite bioinks showing variation in the storage modulus and loss modulus upon commencement of visible light irradiation after 60 seconds. (D) Frequency sweep after photocrosslinking for 300 seconds. (E) Crosslinking rate comparison for various compositions of GelMA-OrnMA hydrogels. (F) Zoomed section highlighting major difference in crosslinking rates.

To investigate further, the photocrosslinking kinetics were assessed using a rheometer in order to identify ink to hydrogel transition and compare various ink compositions. The samples were probed by applying a constant strain of 1% at a constant frequency of 1 Hz and the respective storage modulus (G’) and loss modulus (G”) were recorded. Figure 2C illustrates the photocrosslinking kinetics of various compositions of GelMA and OrnMA composite inks analyzing G’ and G’’ over time. As shown in Figure 2C, prior to visible light exposure at 60 seconds, both G’ and G’’ were low across all ink samples, representing their uncrosslinked states. The visible light irradiation was turned on after 60 seconds, triggering the photocrosslinking cascade and resulting in an immediate increase in G” for each composition. Ink compositions with higher concentrations of OrnMA, such as 10% and 15% formulations exhibited a rapid increase as compared to pristine 5% GelMA and 5% GelMA-5% OrnMA compositions. Among 5% GelMA and 5% GelMA-5% OrnMA compositions, the addition of OrnMA resulted in slightly faster photocrosslinking behavior. This suggests more efficient crosslinking kinetics with increased OrnMA content. With continued light exposure, G’ continued to increase for all samples, indicating successful crosslinking and gel stiffening. The G’ initially increased but then plateaued or decreased, marking a shift from a viscous to a more elastic, solid-like state. However, in the case of 5% GelMA-15% OrnMA ink, the G’ increased rapidly at the onset of light exposure and then decreased after achieving a high value of G’ around 80-90 seconds. This observation can be attributed to a high OrnMA to GelMA ratio of 3:1 in the ink composition. Since both GelMA and OrnMA contain methacryloyl function groups and are uniformly mixed in the ink, the availability of OrnMA is higher at all locations within the ink. Therefore, when the photocrosslinking cascade commences, OrnMA can also react with surrounding OrnMA molecules to form clusters. High availability of OrnMA supports this process and, as a result, the GelMA-OrnMA composite hydrogel contains localized clusters of OrnMA within the matrix. Such a structure can exhibit high brittleness and can easily breakdown under agitation during the rheology measurements resulting in decline in G’.

The light exposure continued for 5 minutes to provide sufficient duration for photocrosslinking and to form a stable hydrogel. Beyond 5 minutes, the light exposure was turned off and subsequent characterization of the gel stability was performed using Frequency sweep as outlined in section 2.3. Figure 2D shows the frequency sweep measurement of the gels formed post photocrosslinking and presents the variation of G’ and G’’ as a function of angular frequency. The G’ remains higher than G’’ across all frequencies during the frequency sweep test, confirming the solid-like nature of the hydrogels after crosslinking. Hydrogels with OrnMA, particularly the 15% OrnMA formulation, exhibit the lowest G’ values across the frequency range compared to pure GelMA due to the reasons mentioned earlier. The frequency-dependent behavior indicates that, while 5% GelMA-15% OrnMA undergoes faster photocrosslinking, its stiffness across various frequencies is lower than other hydrogels. In contrast, 5% GelMA shows the highest G’ values, indicating it retains more stiffness across different angular frequencies. The 5% GelMA-10% OrnMA and 5% GelMA-5% OrnMA formulations fall in between, with moderate G’ values, reflecting a balance between elasticity and stiffness. This behavior can be attributed to the potential plasticizing effect of OrnMA at higher concentrations. While OrnMA promotes faster crosslinking, as observed in Figure 2C with the sharp increase in G’, higher concentrations may introduce a more heterogeneous network, leading to lower overall stiffness (lower G’) at different frequencies. Additionally, a less densely packed crosslinking network may result from the increased OrnMA concentration, which could further explain the reduced stiffness at low frequencies, where the material has more time to respond elastically. Overall, the addition of OrnMA to GelMA significantly enhances photocrosslinking efficiency. The 5% GelMA-15% OrnMA hydrogel forms the weakest network due to induced brittleness from smaller OrnMA chains, however, it shows the fastest crosslinking kinetics, while pure GelMA exhibits slower crosslinking kinetics and higher final stiffness. This behavior of OrnMA may be explained as in the initial stages of gelation due to its role in enhancing cross-linking rate OrnMA forms a more homogeneous network. However, at higher concentrations such as 15 % OrnMA, the hydrogel network might become more heterogeneous and brittle, as the flexibility introduced by OrnMA may disrupt the overall uniformity of the gel structure. This makes the GelMA-OrnMA hydrogels highly applicable in fields requiring tailored stiffness and rapid gelation.

### 3.3. Physical and Microstructure characterization of GelMA-ornithine hydrogels

Next, the physical characteristics of the obtained hydrogels with various compositions of GelMA and OrnMA were examined. First, the microscale structure of the hydrogels was visualized using scanning electron microscopy (SEM). The SEM micrographs in Figure 3A show the cross-sectional porous morphology of the lyophilized hydrogels. The pristine GelMA hydrogels had a consistent porous structure comprising pores of different sizes. Further, the hydrogel pore walls were smooth in appearance. The hydrogel with 5% GelMA-5% OrnMA composition exhibited irregularity in the porous structure with varied pore sizes. Closer examination of the hydrogel pore walls shows a reinforced structure with OrnMA fibrils attached to the hydrogel walls during photocrosslinking. As the OrnMA concentration increased to 10%, the OrnMA fibrils began to form dense clusters. The porous structure of the hydrogel also changed significantly as smaller pores diminished and larger pores became prominent. The clusters of OrnMA fibrils also had distinct tubular morphology. This observation continued as the OrnMA concentration was increased to 15%. Addition of more OrnMA resulted in formation of a greater number of clusters. As described earlier, the availability of larger number of OrnMA molecules in a localized volume results in polymerization of OrnMA first, followed by crosslinking with GelMA macromolecules. The formation of such clusters also supports the argument that higher ratio of OrnMA to GelMA results in increasing granular nature of the hydrogels making them brittle. Increased granularity resulted in brittleness of the hydrogels as observed during photocrosslinking characterization (Figure 2C-D). Presence of such tubular structures and clusters formed by OrnMA within the hydrogels can lead to the formation of a heterogenous microenvironment and provide physical stimulation to the encapsulated cells.

**Figure 3.**
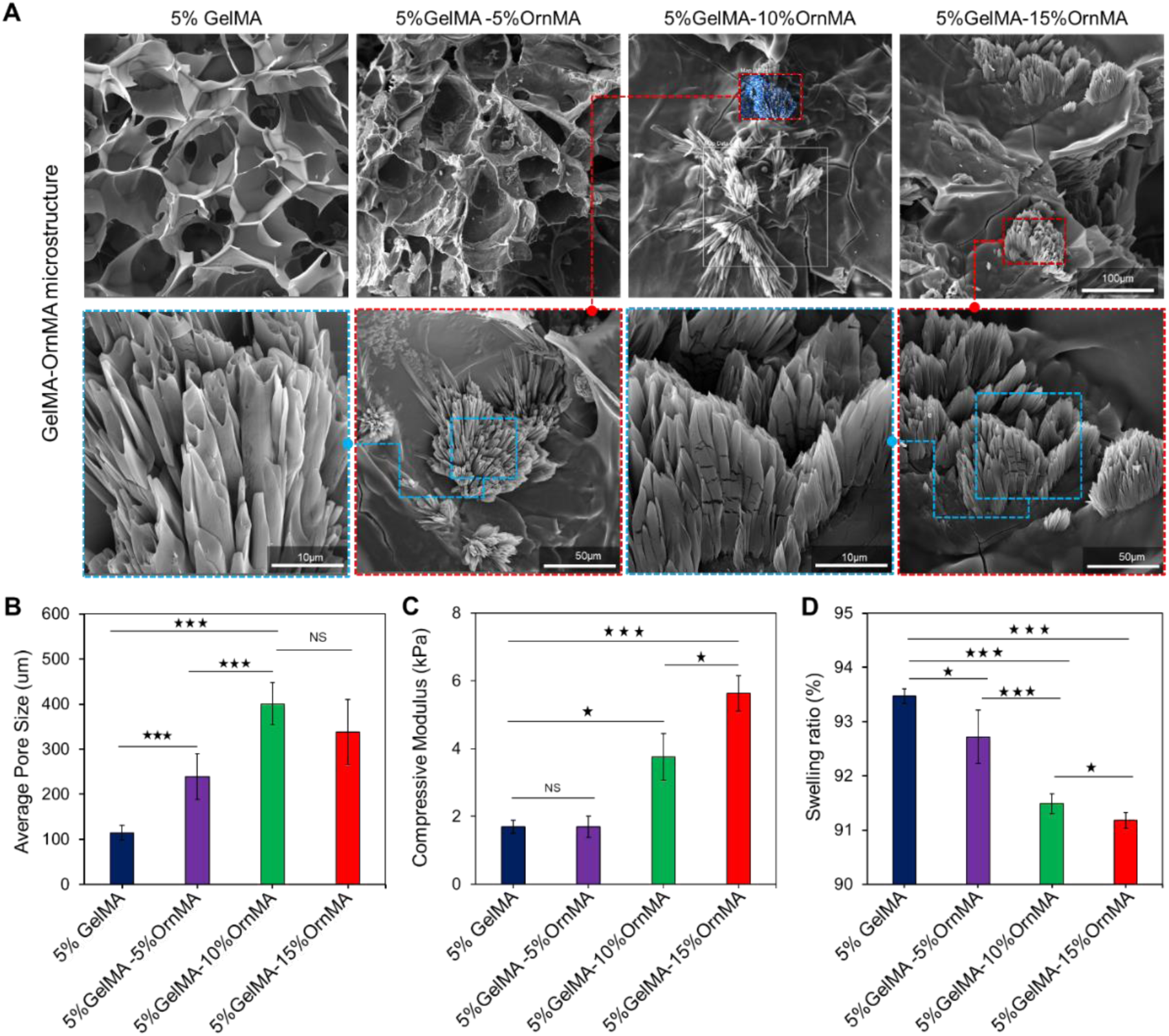
Physical characterization of GelMA-OrnMA composite hydrogels. (A) SEM micrographs showing changes in porous microstructure of of GelMA-OrnMA hydrogels with different concentrations of OrnMA (scale: 100µm). (Insets) OrnMA tubules and clusters were observed within the GelMA walls (scale: 10µm, 50 µm). (B) Average pore size estimated from SEM micrographs of GelMA-OrnMA composite hydrogels (n = 20). (C) Compressive modulus evaluated from unconfined compression of the composite hydrogel hydrogels (n = 5). (D) Water uptake capacity of the hydrogels is represented as swelling ratio hydrogels (n = 5). (NS p > 0.05, *p < 0.05, **p < 0.01, ***p < 0.001, and **** p < 0.0001).

The SEM micrographs were further examined using a custom-built MATLAB script to analyze porous microstructure. As shown in Figure 3B, the average pore size increased to 239.0±50.18 µm, approximately 2 times, by adding 5% OrnMA as compared to the average pore size of 114.68±16.30 µm in pristine 5% GelMA hydrogels. Addition of 10% OrnMA resulted in approximately 3.5 times increase in the average pore size to 400.38±46.46 µm as compared to pristine 5% GelMA hydrogels. Continued addition of OrnMA to a concentration of 15% resulted in slightly lowered average pore size of 338.43±72.22 µm as compared to 10% OrnMA, however, this change was not significant. Although, the average pore size increased with increasing OrnMA concentration, as observed in the SEM micrographs, the overall number of pores declined with OrnMA addition. The hydrogels were further characterized to assess their stiffness as it is an essential property of the microenvironment, and it can provide various biomechanical cues for guiding cell behavior. The hydrogels underwent unconfined compression to obtain the force-displacement data which was used to compute the compressive modulus of the hydrogels. Figure 3C shows that addition of 5% OrnMA did not have any significant influence on the stiffness of the composite hydrogel as compared to pristine 5% GelMA hydrogels and both of these compositions exhibited a compressive modulus of 1.69±0.31 kPa and 1.68±0.27 kPa. However, with addition of 10% and 15% OrnMA, the stiffness of the composite hydrogels increased by two-folds and three-folds, respectively, to 3.75±0.65 kPa and 5.63±0.52 kPa. Despite variations, these hydrogels are comparable to the stiffness range of brain tissue (1-3.5 kPA)^51,52^ with possibility of further tuning their properties. This makes the GelMA-OrnMA hydrogels potentially suitable candidates for recreating neural tissue microenvironment. Beyond porous microstructure and stiffness of the hydrogels, water uptake capacity of the hydrogels is an important property which can influence the microenvironment experienced by the encapsulated cells. In this work, the water uptake capacity is denoted by the swelling ratio described as the ratio of hydrated hydrogel weight to dried hydrogel weight. The water uptake capacity of the hydrogels typically exhibits an inverse trend as compared to the stiffness of the hydrogels^42,45,53^. GelMA-OrnMA followed this observation and exhibited a decreasing trend in hydrogel water update capacity with increasing OrnMA concentrations. The pristine 5% GelMA had the highest swelling ratio of 93.47±0.13% followed by 5% GelMA-5% OrnMA hydrogels at 92.72±0.53%. Further addition of 10% and 15% OrnMA further brought down the swelling ratio to 91.48±0.12% and 91.17±0.22% respectively. Although the water uptake capacity of the GelMA-OrnMA hydrogels decreased with increasing OrnMA concentrations, the swelling ratio remained above 90%. High water uptake capacity of the hydrogels is a desirable property and aids in biocompatibility, thus, the GelMA-OrnMA hydrogels are expected to exhibit high biocompatibility.

### 3.4. Electrical characterization of GelMA-OrnMA hydrogels

Hydrogels based on GelMA are conventionally non-conductive, limiting the application of GelMA hydrogels for culturing and stimulating electrically responsive cell types, such as neurons, astrocytes, and cardiomyocytes. Therefore, we introduced OrnMA to form nanocomposite hydrogels with GelMA and covalently bind zwitterionic functional groups to the hydrogel matrix. We examined the electrical conductivity of pristine GelMA and GelMA-OrnMA hydrogels using distilled water (DW) and PBS as solvents. PBS was used to mimic the physiological and cell culture salinity environment, as shown in Figure 4. First, a custom sample platform, as shown in Figure 4A, was constructed to consistently hold the samples during the measurements as sample movements may introduce error in measurements. After measuring the resistance of the samples, the hydrogel dimensions were used to compute electrical conductivity of the pristine GelMA and GelMA-OrnMA composite hydrogels. When distilled water was used as solvent to prepare the hydrogels, pristine GelMA hydrogels exhibited low conductivity of 1.06±0.32 mS/m due to their intrinsic non-conducting nature^54,55^. With the addition of 5% OrnMA, the conductivity of the hydrogels increased significantly to 13.08±1.67 mS/m. When the OrnMA concentration was increased further to 10% and 15%, the conductivity steadily increased to 26.68±2.76 mS/m and 32.85±3.24 mS/m respectively. This emphasized that addition of OrnMA introduced components in the hydrogels supporting for electrical conductivity. On the other hand, when the PBS was used as a solvent, the resulting GelMA hydrogels exhibited a conductivity of 4.35±0.65 mS/m. Upon addition of 5% OrnMA, a striking change in the conductivity which increased approximately 6.5 times to 28.34±6.28 mS/m. Hydrogels with 10% OrnMA showed further increase in the conductivity to 49.93±5.03 mS/m. Addition of 15% OrnMA resulted in even higher conductivity of 81.72±17.30 mS/m which was approximately 18 times the conductivity of pristine GelMA hydrogels. Although, the addition of OrnMA resulted in increased conductivity of the hydrogels, in presence of PBS as solvent, the resulting conductivity was strikingly higher. This emphasized on potential contribution of ionic species in PBS in the conduction mechanism. A possible conduction mechanism can be described by considering the structure of the hydrogels formed as a result of photocrosslinking. Figure 4C shows the photocrosslinked structure of the pristine GelMA hydrogels with ionic species distributed throughout the matric. In contrast, the photocrosslinking of GelMA-OrnMA inks result in a matrix with covalently bound zwitterionic groups (Figure 4D). These zwitterionic groups influence the ionic species in matrix and provide them with hopping sites to facilitate ionic conduction.

**Figure 4.**
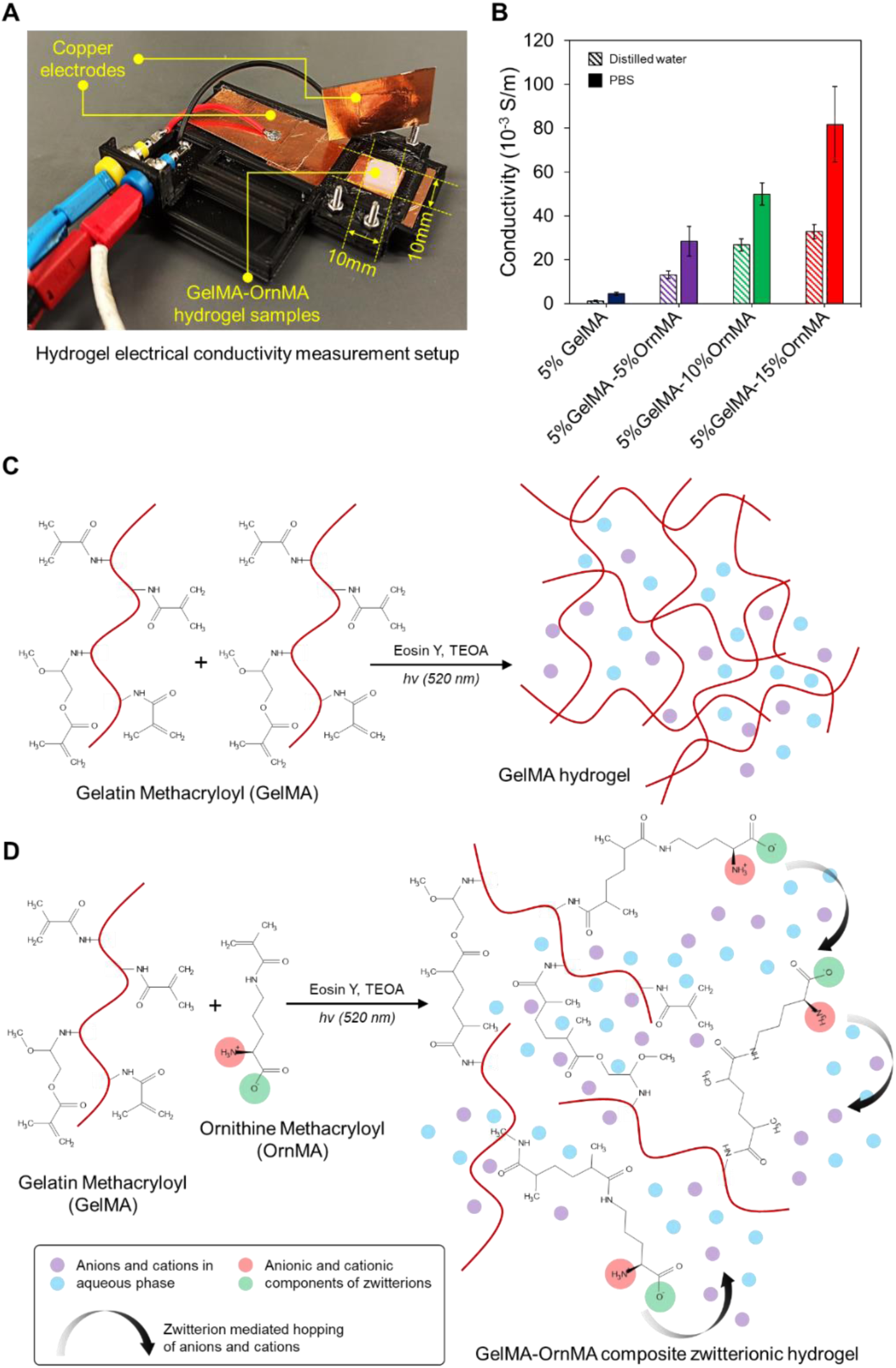
(A) Electrical conductivity measurement setup. (B) Electrical conductivity results for GelMA-OrnMA hydrogels. (C) Photocrosslinking mechanism of GelMA using Eosin Y and TEOA. (D) Anticipated conductivity mechanism for GelMA-OrnMA hydrogels.

The electrical conductivity achieved with GelMA-OrnMA hydrogels matched and surpassed the conductivity exhibited by some of the other zwitterionic hydrogels^56,57^. The GelMA-OrnMA hydrogels also exhibited the required conductivity levels for the stimulation of responsive cell types. As the content of OrnMA was increased up to 10%, the conductivity of hydrogel exceeded 10–2 S/m, which satisfied the needs of neuron cells for micro-current stimulation ^58–60^. The variation in conductivity across all hydrogels exhibited a convex-up behavior with increasing OrnMA concentration when distilled water was used as the solvent as presented in Figure 4B. Similarly, when PBS was used as solvent, although the conductivity increased with OrnMA concentration, the change in conductivity declined. This implies that the electrical conductivity of the composite hydrogels does not increase linearly with OrnMA concentration and is expected to saturate at high OrnMA to GelMA ratio. However, at OrnMA to GelMA ratio is increased, the hydrogels develop a granular nature, and the stability of the hydrogels is compromised as described earlier. Therefore, GelMA-OrnMA composite hydrogels up to 15% OrnMA and 5% GelMA deem well suited for engineering tissues requiring electrical conductivity and mechanical stability.

### 3.5. Biocompatibility assessment

The biocompatibility study results depicted in Figure 5 examine the effect of different concentrations of GelMA-OrnMA hydrogels on the encapsulated astrocytes, over time (days 1, 3, 4, and 5). The study assesses both cell viability and morphology as well as the proliferation of these cells within the GelMA-OrnMA matrix as per the procedure outlined in section 2.7. Figure 5A shows live/dead stained images portraying the viability of astrocytes encapsulated in the GelMA-OrnMA hydrogel matrix from day 1 to day 5. Green fluorescence indicates live cells stained with Calcein-AM, while red fluorescence denotes dead cells stained with Propidium iodide. In all GelMA-OrnMA compositions, there is a greater number of live cells, which suggests high cell viability throughout the varying GelMA-OrnMA matrices. The increasing density of cells from day 1 to day 5 indicates that the GelMA-OrnMA hydrogel matrix provides a suitable environment for cell survival. With increasing culture time from day 1 to day 4, the number of live cells appears to increase indicating that the GelMA-OrnMA hydrogels promote both the viability and potential proliferation of the astrocyte. Notably, the live/dead staining shows fewer dead cells (red) across all the samples, emphasizing the biocompatibility of the hydrogels.

**Figure 5.**
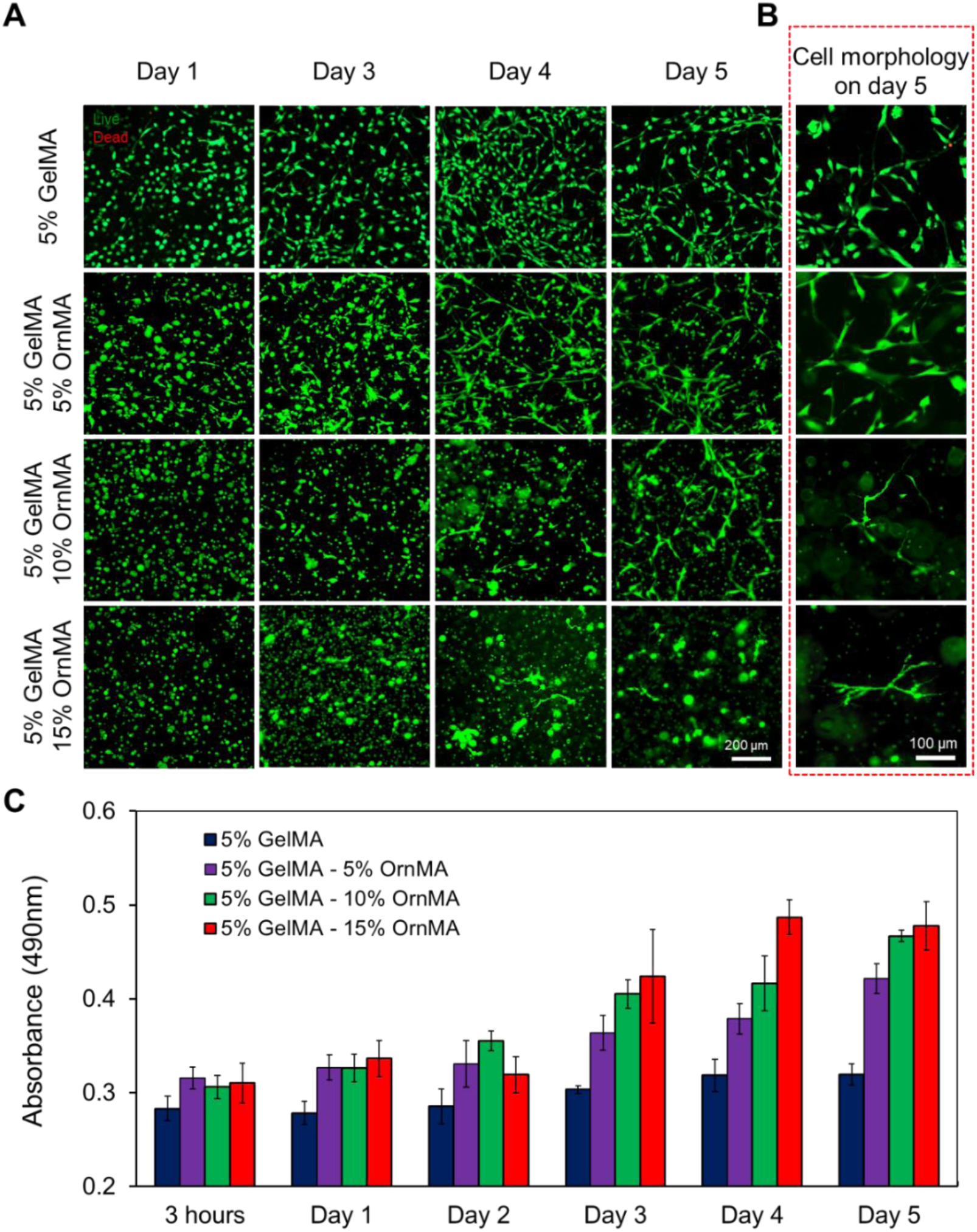
Biocompatibility study of GelMA-OrnMA composite hydrogels. (A) Live/Dead staining of 3D encapsulated astrocytes from day 1 to day 5 within the different concentrations of GelMA-OrnMA matrix, (B) Variations in astrocytes morphology on day 5 within the different concentrations of GelMA-OrnMA matrix, (C) Proliferation of encapsulated cells within various GelMA-OrnMA matrices examined using XTT proliferation assay.

Figure 5B shows the morphology of astrocytes at day 5 in the hydrogel matrix with different GelMA-OrnMA concentrations. Astrocytes exhibit different morphologies based on their interaction with the extracellular matrix. The elongated and dendritic morphology of astrocytes is an indication of how well the GelMA-OrnMA matrix mimics the natural environment of the cells. The images also reveal an increasing complexity in astrocyte processes as the concentration of GelMA-OrnMA increases. Further, in Figure 5B, the astrocytes display a more organized network, reflecting that the GelMA-OrnMA concentration is influencing astrocyte interactions and potentially their functionality. Such an arrangement is crucial for their physiological role, and this suggests that GelMA-OrnMA matrices can support not just cell survival but also the physiological functions of encapsulated astrocytes.

Figure 5C shows the results of an XTT proliferation assay, which quantitatively measures cell proliferation over time. It shows that the astrocytes proliferate within different concentrations of the GelMA-OrnMA hydrogel over 5 days. Across all concentrations, there is an increasing trend in proliferation, indicating that astrocytes proliferate well within the GelMA-OrnMA matrix. Notably, the highest proliferation rates are observed in the hydrogel matrices with a concentration of 10-15% OrnMA. On the other hand, the control sample (5 % GelMA) shows relatively slower proliferation, though still significant compared to day 1, indicating that the OrnMA concentration plays a critical role in cellular proliferation of the astrocytes. These results show that GelMA-OrnMA composite hydrogels are highly biocompatible and support the survival, morphology, and proliferation of encapsulated astrocytes. The Live/Dead staining demonstrates the hydrogels’ ability to maintain high cell viability over time, with very few dead cells observed, while the proliferation assay confirms that astrocytes can actively divide within the matrix, especially at higher OrnMA concentrations. The hypothesis behind it is 1) increasing the concentration of ornithine, increases the stiffness of the hydrogel which affects the cell cycle regulations and cytoskeletal remodeling. 2) Increasing the concentration of ornithine increases the electrical conductivity of the hydrogel matrix which enhances cell signalling and enhance in return cell-to-cell communication and astrocytes differentiation and proliferation^61^. Earlier studies have highlighted that the mechanical strength and the functionality of the astrocytes could be enhanced by increasing the stiffness of the microenvironment^62^. In literature, healthy brain stiffness is around 0.1-1 kPa which is relevant to our control pristine GelMA samples without OrnMA (Figure 3C), and meningioma is 4 kPa. A. Bizanti et al.^62^ studied the increase of intercellular stresses and strain energy due to the increased substrate stiffness. Similar studies mentioned the mechanosensing of astrocytes to changes in hydrogel stiffness^63^.

### 3.6. Engineered neural tissues

To study astrocytes morphology, an in-vitro experiment included astrocytes encapsulated inside the crosslinked samples forming GelMA-OrnMA 3D matrices, and F-actin/DAPI staining was performed to observe cell growth. Figure 6A shows the encapsulation process of astrocytes within the hydrogel. Figure 6B shows that the cells proliferated and adhered to the matrix in all hydrogel compositions when maintained 5 days in culture. Further, it was also noticed that the astrocytes had different organization patters according to the OrnMA concentration in the matrices. This might be because of the electrical conductivity of zwitterionic OrnMA which could influence the cell growth and the directionality of the cell’s elongations. The astrocytes encapsulated in pristine GelMA matrix did not exhibit any specific orientation and were randomly organized throughout the matrix. With introduction of OrnMA, an increased population of astrocytes was observed in all GelMA-OrnMA hydrogels as compared to pristine GelMA hydrogels. Furthermore, 5% GelMA-5% OrnMA samples exhibited the cells’ tendency to form an interconnected network. With 10% OrnMA, the astrocyte network became denser, and a pseudo-aligned organization was evident. The 10% OrnMA matrices also exhibited relatively more elongated morphology of the cells. With 15% OrnMA in hydrogels, a similar network forming tendency was observed along with thin stretched processes. However, the cell density was lower as compared to 10% OrnMA samples. This can be attributed to the increased stiffness of the 15% OrnMA matrices. The nuclei and F-actin stained images were further processed to extract two features of interest – (i) eccentricity of nuclei as shown in Figure 6C and (ii) cell area as shown in Figure 6D. Both of these features can describe the morphology of the cells in the 3D matrices. As shown in Figure 6C, the eccentricity of the nuclei shifted from 0 (uniformly spread cell morphology) towards 1 (highly elongated cell morphology) with addition of OrnMA. The pristine GelMA matrix demonstrated a mixed behavior from the encapsulated cells where some cells exhibited lower eccentricity value and some exhibited high eccentricity. As the concentration of OrnMA was increased to 15%, majority of the cells exhibited eccentricity value close to 1 implying an elongated morphology similar to the *in vivo* microenvironment. This observation was further supported by the measurement of cell area shown in Figure 6D. Cells with uniformly spread morphology exhibit larger cell area in the images. The cells in 5% GelMA samples had an average cell area of 0.043±0.015 µm^2^ which steadily decreased to 0.020±0.002 µm^2^, 0.014±0.004 µm^2^ and 0.003±0.002 µm^2^ in matrices with 5%, 10% and 15% OrnMA respectively. This change can be a result of increasing electrical conductivity of the microenvironment which potentially improves the chemokine signaling between farther cells and induces the cells to form interconnects across longer distances. Furthermore, formation of such interconnects can lead to a better replication of the astrocyte networks in native tissues and should be investigated in detail in co-culture models including other neural cell types. The striking changes in the morphology of the astrocytes and their organization closer to *in vivo* microenvironment suggest GelMA-OrnMA hydrogels to be suitable candidates for developing engineered neural tissues for replicating the complexity of native tissue more accurately.

**Figure 6.**
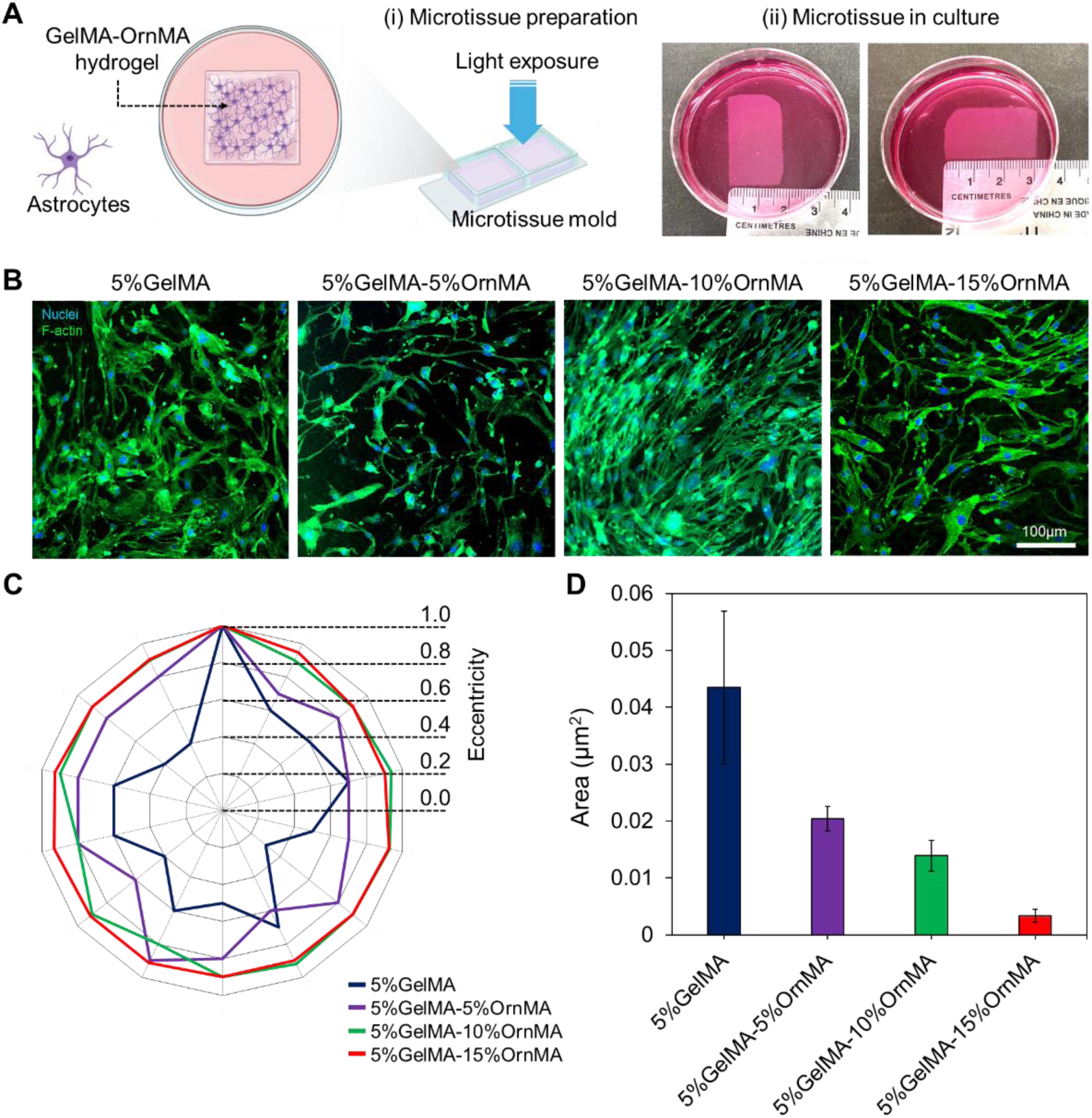
(A) 3D Encapsulation of astrocytes within GelMA-OrnMA hydrogels (i) demonstrated by schematic and (ii) images of cell-encapsulating hydrogels. (B) Fluorescent images with stained nuclei and cytoskeleton showing organization and morphology of U118 cells in hydrogels with different OrnMA concentration. (C) Eccentricity of the nuclei measured from the fluorescence images. (D) Comparison of cell area measured in images corresponding to various GelMA-OrnMA composite hydrogels.

Beyond studying the interaction of astrocytes with GelMA-OrnMA hydrogels, these bioinks were also explored for biofabrication. A high-resolution structure was bioprinted (Figure 7) to demonstrate the printing capability using visible light. This was feasible due to the addition of OrnMA as it increases the photocrosslinking rate as well as the mechanical stability of the resulting hydrogels. Figure 7A shows the designed scaffold along with the dimensions. Figure 7B illustrates the SLA-printed scaffold colored with red dye to enhance visualization. Next, Figure 7C shows the brightfield image of the bioprinted structure with astrocytes on days 0, 3, and 4 showing the progression of cell growth as well as distortion of microscale features in the hydrogels due to matrix remodeling. After day 3, the initial elongation of astrocytes can be observed in the bioprinted structure (marked with red arrows). Further, a greater number of astrocyte elongations and interconnections were observed after day 4. After maintaining the samples for 5 days in culture, in order to examine the cellular morphology, we stained the bioprinted structures using DAPI/Phalloidin. Figure 7D shows the morphology of astrocytes with F-actin (green: Phalloidin) and nuclei (blue: DAPI). It can be seen that the majority of the cells have elongated and formed an interconnecting network characteristic of astrocytes growth *in vivo*.

**Figure 7.**
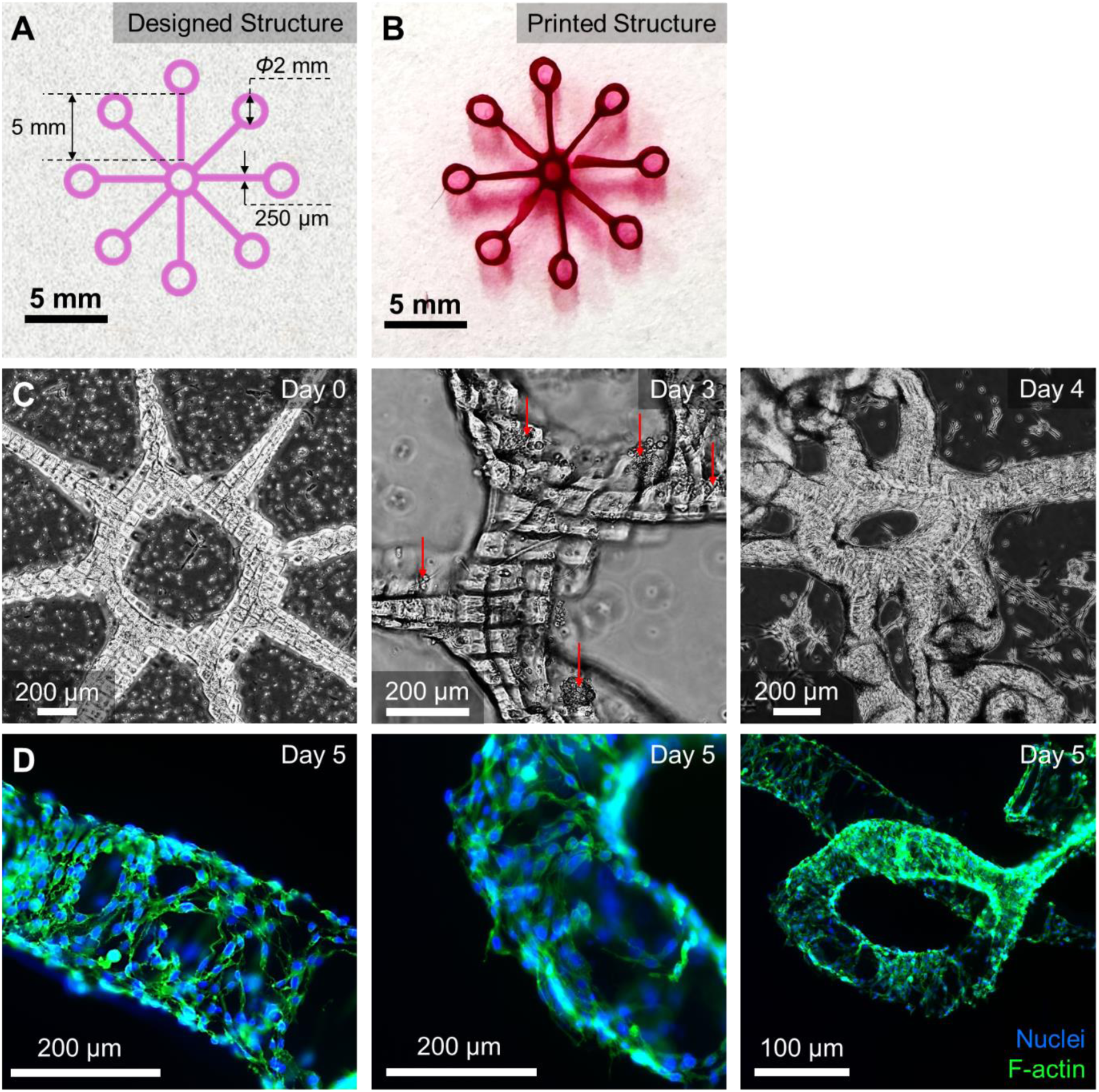
Bioprinting high-resolution structures. (A) Design of a star shape with ring ends bioprinted to demonstrate high-resolution fabrication. (B) Fabricated structure using visible light stereolithography bioprinting (added color to enhance visualization). (C) Brightfield images showing the encapsulated cells in various parts of the structure at day 0, 3 and 4. (D) Fluorescent images showing the organization and morphology of the encapsulated cells in the straight branches and rings at day 5.

## 4. Conclusions

This study demonstrated the development of a novel ornithine-GelMA bioink that can be used for tissue engineering applications. In this study, the physical characteristics of the composite hydrogel were investigated. The morphological study using SEM demonstrated the presence of ornithine within the walls of the cross-linked GelMA matrix. The reinforcement of GelMA walls with ornithine resulted in enhancing the stiffness of the hydrogel threefold by increasing the concentration of ornithine. In addition, the main advantage of using ornithine is to maintain the transparency of the hydrogel. As noticed in the transparency data, the transparency of the hydrogel didn’t change by adding 5-15% ornithine to pristine GelMA. Moreover, the electrical conductivity of the hydrogel was tested and the presence of the free ions from the ornithine reaction with the GelMA chains resulted in the ionic conductivity under applied bias. The electrical conductivity of the hydrogel demonstrated an enhancement while increasing the concentration of ornithine. The increase of ornithine concentration resulted in increasing the number of free ions which increases the electrical conductivity of the hydrogel.

Overall, the effect of ornithine was observed as a newly developed bioink for bioprinting and tissue engineering applications. It is noticed that there is a change in the morphological, physical, and biological characteristics of pristine GelMA in the presence of ornithine. Ornithine, an amino acid, is made in the body and can be integrated with the GelMA chains and form a novel GelMA-OrnMA hydrogel which can be used in bioprinting applications. This article discusses the effect of 0, 5, 10, and 15 % ornithine mixed with 5% GelMA in the change of microstructure and physical properties of GelMA. OrnMA resulted in stiffening the walls of GelMA and changing the morphology of the walls. It is noticed that the compressive modulus increased 4 times with 15 % ornithine. The swelling ratio decreased by increasing ornithine concentration, because of the change in the pore size of the hydrogel. In addition, it is noticed that GelMA-OrnMA composite bioink were biocompatible, proven by 3D cell viability experiments with astrocytes.

## Acknowledgement

This work was supported by the Natural Sciences and Engineering Research Council of Canada (NSERC) Discovery Grants and Canada Foundation for Innovation John R. Evans Leaders Opportunity Fund. Kartikeya gratefully acknowledges Alberta Innovates for providing financial support through the Postdoctoral Fellowship.

## Data availability

All data that support the findings of this study are included within the article (and any supplementary files).

